# Flow Currents Support Simple and Versatile Trail-Tracking Strategies

**DOI:** 10.1101/2023.12.15.571932

**Authors:** Haotian Hang, Yusheng Jiao, Josh Merel, Eva Kanso

## Abstract

Aquatic animals offer compelling evidence that flow sensing alone, without vision, is sufficient to guide a swimming organism to the source of an unsteady hydrodynamic trail. However, the sensory feedback strategies that allow these remarkable trail tracking abilities remain opaque. Here, by integrating mechanistic flow simulations with reinforcement learning techniques, we discovered two simple and equally effective strategies for hydrodynamic trail following. Though not *a priori* obvious, these strategies possess parsimonious interpretations, analogous to Braitenberg’s simplest vehicles, where the agent senses local flow signals and turns away from or toward the direction of stronger signals. A rigorous stability analysis shows that the effectiveness of these strategies in robustly tracking flow currents is independent of the type of sensor but depends on sensor placement and the traveling nature of the flow signal. Importantly, these results inform a suite of versatile strategies for hydrodynamic trail following applicable to both vortical and turbulent flows. These insights support the future design and implementation of adaptive real-time sensory feedback strategies for autonomous robots in dynamic flow environments.

## Introduction

The physical environment plays a pivotal role in shaping the sensory, behavioral and survival strategies of living organisms [1–4]. Chemotactic bacteria inhabit a viscous flow environment, where chemical concentrations are governed by diffusion [5, 6] (Fig. 1A), and employ a run-and-tumble strategy to navigate toward nutrient sources and away from toxins [7–9]. Minute insects and crustaceans, including drosophilas [10–12], moths [13] and copepods [14–17], inhabit a turbulent flow environment. To find potential mates, they detect pheromones carried by turbulent wind or water current, where encounters with odor patches are random and intermittent [18] (Fig. 1B). Under such information-scarce conditions, trail-following requires elaborate sensory and movement strategies that rely on constructing spatial maps [18] or remembering flow direction and temporal history of the detected signal [19]. Larger animals, such as fish [20–22] and harbor seals [23, 24], do not experience the world as merely laminar or turbulent. They encounter a rich array of flow patterns and coherent vortex structures [25–27] created by conspecifics, predators, prey, and even abiotic sources (Fig. 1C). These flows contain, in principle, useful information about the source generating them that can be deciphered by an aspiring follower [26–28]. However, effective feedback strategies for tracking coherent vortical flows remain largely unexplored.

**Figure 1:**
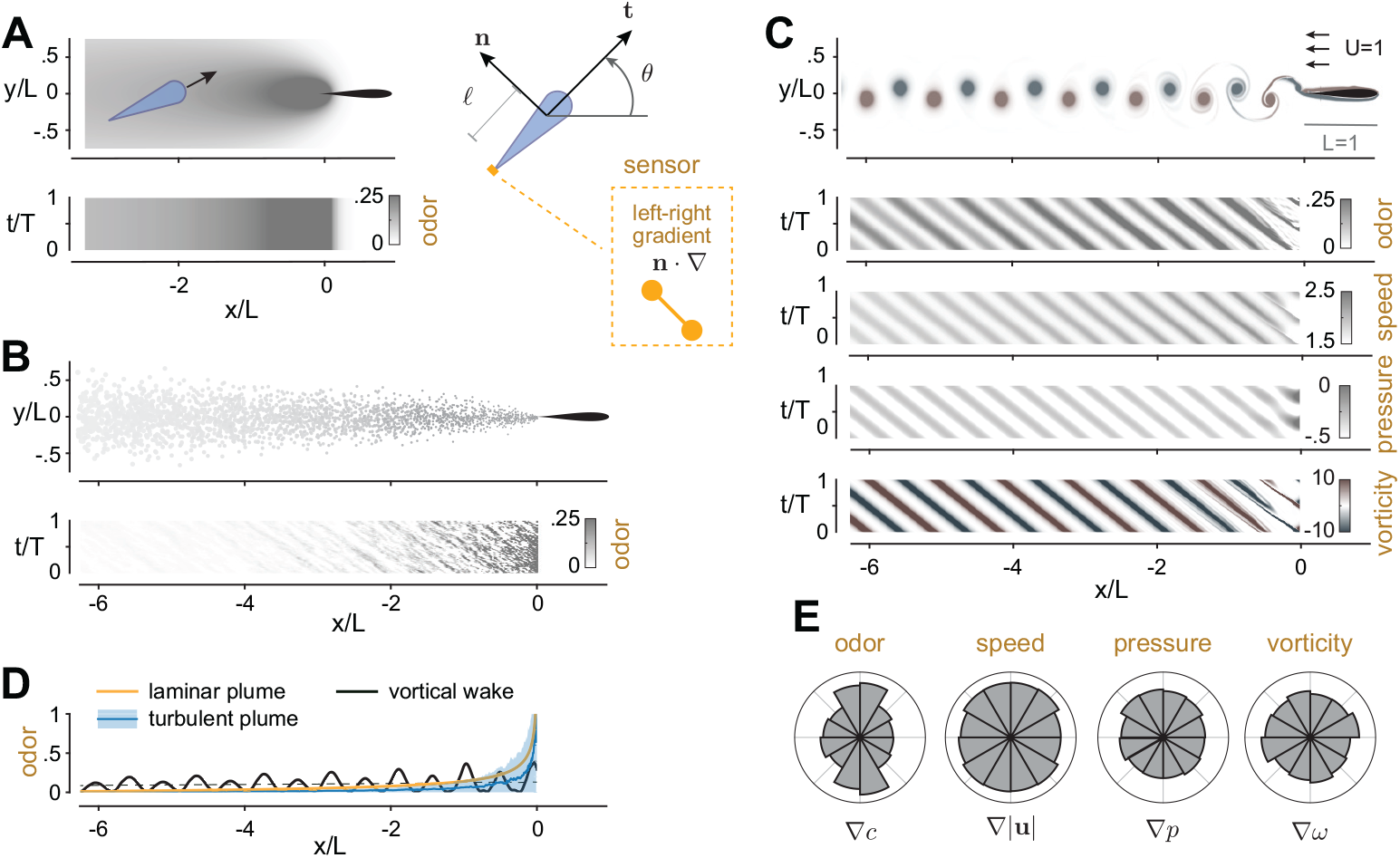
Scented and odorless trails at different scales. **A**. In laminar plumes, odor is released from a source at a fixed emission rate *R* = 1 and is advected by a uniform background flow while undergoing molecular diffusion at Pe = 10: (top row) steady-state solution, (bottom row) concentration along the midline *y/L* = 0. An agent unaware of the source location is modeled as a swimmer with body-fixed frame (**t, n**), moving at a constant speed *V* and with control over its rate of orientation 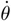, where *θ* is the angle between its heading direction **t** and the inertial *x*-axis. **B**. To simulate turbulent plumes, odor packets are released stochastically from the source and transported by the turbulent wind: (top row) odor packets of decaying strength undergoing random cross-stream perturbations, (bottom row) odor concentration along the midline *y/L* = 0. **C**. Hydrodynamic trail created by a Joukowsky foil undergoing pitching oscillations in uniform background flow *U* = 1 with tailbeat amplitude *A* = 0.2 and frequency *f* = 1.25, at Re = 5000 and St = 0.25. Odor released from the airfoil’s trailing edge is carried by the flow field and undergoes diffusion at Pe = 10, 000. (bottom rows) Physical quantities along the midline *y/L* = 0, including odor, flow speed, pressure, and vorticity, exhibit traveling wave characteristics. **D**. Decay rates of the concentration fields in panels A, B, C along the midline *y/L* = 0: vortical flow shows the weakest decay. **E**. Statistics of gradient directions of each of the physical quantities – odor, speed, pressure, and vorticity – evaluated along a regular grid in the flow field of panel C. Gradients have no preference for upstream or downstream directions.

Hydrodynamic trail following is a critical task for many aquatic organisms [20, 24], with direct implications for developing navigation strategies in underwater robotic vehicles [29–32]. Various organisms detect flow signals [20–22] to negotiate conditions with reduced or no visibility [33] or to reduce the cost of locomotion [34–36]. The flow-sensitive mechanoreceptors vary by species, from the lateral line system in fish [37–41] to the whiskers of harbor seals [24, 42]. The underlying principles that allow flow detection [43, 44] have direct implications to designing bio-inspired flow sensors [45–47]. Here, we are concerned, not with sensor design, but with the navigation ability of flow-sensitive swimmers. Indeed, fish [20, 22, 33] and harbor seals [24, 42] follow hydrodynamic trails for underwater navigation not only over short distances, as in the final stage of mate or prey localization [33], but also to track and locate distant objects over considerable length and time spans [20, 21, 24, 42]. Additionally, olfaction is often integrated with flow sensing [22] as seen in the homing behavior of the Pacific salmon [48]. Yet, the minimal sensory feedback laws required for a swimmer to respond to local flow signals and follow hydrodynamic trails remain obscure.

Early efforts for designing control strategies to navigate water currents date back to the work of Zermelo [49], where a swimmer is required to navigate toward a target while being influenced by a well-defined laminar flow field. More recently, techniques from optimal control [50, 51], model predictive control [52] and reinforcement learning [50, 51, 53] were applied to solve this problem in unsteady flow fields, including turbulent [53] and vortical flows [50–52]. Interestingly, local [50] and egocentric [51] flow sensing abilities are sufficient to enable the agent to perform efficient navigation, with performance comparable to time-optimal control utilizing full knowledge of the flow field. Behavioral strategies relying only on local sensory information were also devised for soaring birds [54, 55], schooling fish [56], and vertical migration of copepod in turbulent flows [17].

Hydrodynamic trail tracking is a related yet different problem. Instead of moving toward a target whose location is independent of the flow source, the navigator is tasked to move toward the source that generates the flow. This navigation problem is often considered in the context of tracking turbulent plumes, where odor patches are transported randomly [57, 58]. For example, infotaxis uses Bayesian-based methods to build and update a spatial map of the possible location of the source [18]. Alternative strategies for tracking turbulent plumes rely on reinforcement learning [19, 59] or use a finite-state machine approach with limited memory [60]. But turbulent odor tracking is distinct from tracking hydrodynamic trails characterized by long-lasting coherent flow structures [25, 61, 62]. To track such trails, strategies where the agent collects a time history of the local flow signal and acts accordingly were shown to function successfully in a variety of vortical flows [27, 63]. While promising, these handcrafted strategies are suboptimal in terms of sensory requirements.

In this study, we use deep reinforcement learning to uncover efficient tracking strategies that rely purely on local and instantaneous flow sensing. Specifically, we consider a trail-following task with two main features: (1) the swimmer can only sense local flow information, and (2) the swimmer can only sense gradients of the local flow, not the flow itself. That is, we consider a scenario with a completely blind swimmer with no prior or developed knowledge of the source location, inertial coordinates, or overall flow direction. These stringent constraints pose significant challenges to robustly following hydrodynamic trails, whether by locally sensing the flow itself or by sensing the odor concentration carried by the flow. We thus explore a variety of odor and flow sensors. We find parsimonious strategies for trail following, where success depends on sensor placement and the traveling nature of the flow signal, but is largely independent of the sensor type. Importantly, we demonstrate that these strategies can easily translate from coherent to turbulent flows. To conclude, we compare the strategies developed in this work to existing approaches where the navigator has knowledge of an external reference frame [32, 50], overall flow direction [19], vision-based information [50, 51, 56], or past sensory information [18, 19, 59] (Table 1), and we discuss the implications of these findings for future underwater robotic applications.

**Table 1:**
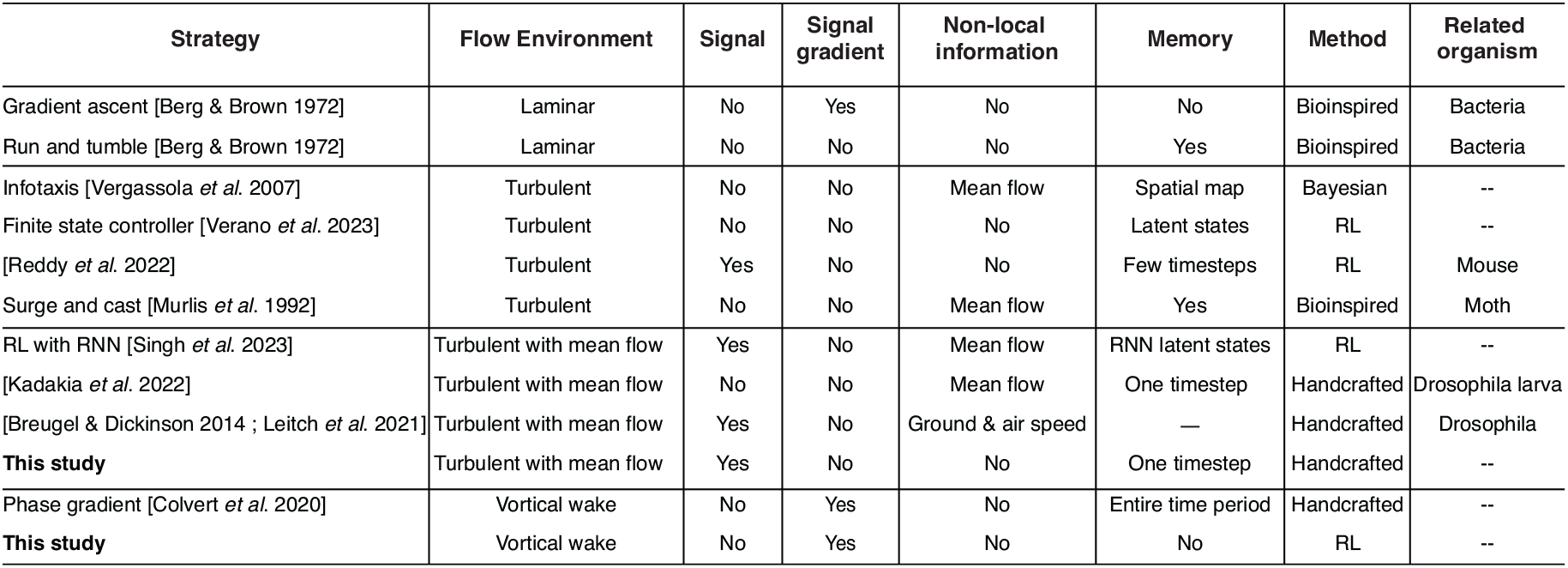
Comparison to existing trail-following strategies. Sensory feedback strategies employed by an autonomous agent for tracking a signal field to its source. We proposed two new strategies that track turbulent plumes and vortical wakes, respectively. In turbulent plumes, our strategy does not require knowledge of the mean flow direction, unlike in [11, 19, 58, 103], and uses less memory and storage than existing strategies designed for the turbulent regime [18, 60, 104]. In vortical flows, our strategies respond instantaneously to the spatial gradient with no memory requirements, unlike in [27], and its performance is optimized using deep reinforcement learning.

| Strategy | Flow Environment | Signal | Signal gradient | Non-local information | Memory | Method | Related organism |
| --- | --- | --- | --- | --- | --- | --- | --- |
| Gradient ascent [Berg & Brown 1972] | Laminar | No | Yes | No | No | Bioinspired | Bacteria |
| Run and tumble [Berg & Brown 1972] | Laminar | No | No | No | Yes | Bioinspired | Bacteria |
| Infotaxis [Vergassola *et al.* 2007] | Turbulent | No | No | Mean flow | Spatial map | Bayesian | -- |
| Finite state controller [Verano *et al.* 2023] | Turbulent | No | No | No | Latent states | RL | -- |
| [Reddy *et al.* 2022] | Turbulent | Yes | No | No | Few timesteps | RL | Mouse |
| Surge and cast [Murlis *et al.* 1992] | Turbulent | No | No | Mean flow | Yes | Bioinspired | Moth |
| RL with RNN [Singh *et al.* 2023] | Turbulent with mean flow | Yes | No | Mean flow | RNN latent states | RL | -- |
| [Kadakia *et al.* 2022] | Turbulent with mean flow | No | No | Mean flow | One timestep | Handcrafted | Drosophila larva |
| [Breugel & Dickinson 2014 ; Leitch *et al.* 2021] | Turbulent with mean flow | Yes | No | Ground & air speed | — | Handcrafted | Drosophila |
| **This study** | Turbulent with mean flow | Yes | No | No | One timestep | Handcrafted | -- |
| Phase gradient [Colvert *et al.* 2020] | Vortical wake | No | Yes | No | Entire time period | Handcrafted | -- |
| **This study** | Vortical wake | No | Yes | No | No | RL | -- |

## Results and Discussion

### Laminar, Turbulent, and Vortical Hydrodynamic Trails

Consider a source of size *L* moving in a fluid at a constant speed *U* with a constant odor emission rate *R*. The characteristics of the odor concentration field depend on the length and speed of the source, as well as the fluid environment, such as water or air. At the micron scale in water, the Reynolds number Re = *UL/ν*, where *ν* is the kinematic viscosity of the fluid, describing the ratio of inertial forces to viscous forces, is nearly zero. The Péclect number Pe = *UL/D*, where *D* is the molecular diffusivity of the odor in fluid, describes the ratio of advective to diffusive time scales and is typically small (less than 100) [5, 6, 64, 65]. Thus, the odor concentration diffuses into a laminar plume, governed by the advection-diffusion equation, with a single maximum at the source location (Fig. 1A, Methods). For a millimeter-scale insect moving in air, Re and Pe are both large. The signal field is transported by a turbulent flow (Fig. 1B, Methods). Here, we employed a particle-based two-dimensional plume model to simulate the random release of odor packets from the source [19, 66]: the odor packets get advected by a mean downstream flow and random cross-stream perturbations, while they decrease in intensity and increase in size over time.

For a source of moderate size and speed, such as a swimming fish [20, 24, 25], the wake is characterized by coherent vortical structures (Fig. 1C, Methods). Here, we solved the Navier-Stokes equation, using an open source computational fluid dynamics (CFD) solver [67–69], to obtain the flow field past a pitching airfoil following sinusoidal oscillations in a uniform background flow, where we set the Reynolds number to Re = 5000 and the Strouhal number St = *f L/U* to St = 0.25, with *f* = 1*/T* being the flapping frequency and *T* the flapping period. This Strouhal number is comparable to those observed in swimming fish, linking the frequency of tailbeat oscillations to the swimming speed and bodylength (Methods). We solved for the concentration field numerically considering the advection-diffusion equation at Péclect number Pe = 10^4^. Considering the odor concentration as the signal field detectable by an aspiring follower, we find that it resembles a traveling wave characterized by wavelength *λ* and frequency *f*. In fact, this traveling wave character emerges in all physical quantities, odor, speed, pressure, and vorticity, in the wake of the source (Fig. 1C).

Whereas the signal field in the laminar plume (Fig. 1A) has a single local maximum at the source, in the turbulent (Fig. 1B) and vortical (Fig. 1C) trails, an aspiring follower encounters multiple local maximum and minimum. Importantly, the signal decays downstream at different rates (Fig. 1D): in the laminar and turbulent regimes, the signal decays exponentially down-stream because of molecular and turbulent diffusion [70], whereas in comparison, the signal lasts much longer in the vortical wake (Fig. 1D). For an aspiring trail follower, the dynamics of the signal fields are intimately connected to which sensing and response strategies are viable. While in the laminar flow regime, it is feasible to measure the gradient of the signal field and use gradient ascent methods to reach the source, spatial gradients are not well defined in the turbulent regime, and gradient ascent tends to get stuck in local maxima in the vortical flow regime. We address these challenges first in the intermediate coherent flow regime, where we develop trail-following strategies that rely on flow or odor sensing. We then discuss extensions of these strategies to the turbulent regime.

### Learning To Track Vortical Trails

We modeled the trail follower (Fig. 1A) as a self-propelled artificial agent, with no inertia, moving at a constant speed *V* and heading at a time-dependent angle *θ*(*t*) from the *x*-axis [71– 73]. We first considered an agent with mechanosensing abilities, where inspired by the fish lateral line system [21, 39, 40] and functional evidence suggesting that flow receptors are optimized to correlate with differential flow signals [74, 75], we considered two sets of flow sensors located at 𝓁_head_ and 𝓁_tail_ from the center, which measure the local gradients of flow speed normal to the agent’s swimming direction *s*_head_ = (**n** · ∇|**u**|)_head_ and *s*_tail_ = (**n** · ∇|**u**|)_tail_. Aside from these two sensory measurements, the agent is at all times ignorant of all other information about the flow field and blind to its position relative to the source generating the flow. To respond to this limited and localized information without memory of past measurements, we provided the agent with control over only its angular velocity 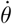, as opposed to direct control over its heading angle *θ* [50]. The agent’s motion is thus described by the nonholonomic unicycle model [71–73],

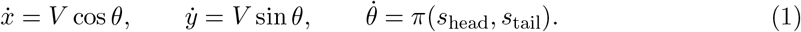

We aimed to learn a policy 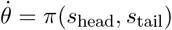 that guides the agent to track the vortical wake to its source from trial-and-error interactions with the flow environment. We employed model-free, deep reinforcement learning based on the Proximal Policy Optimization algorithms [76, 77] (Methods). Actions taken by the policy are restricted to the range of 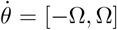 to avoid unrealistically sharp turns. Each training session consisted of 30,000 episodes, with parameter values Ω = 3 *U/L, V* = 0.25*U*, and 𝓁_head_ = 𝓁_tail_ = 0.25*L*. In each episode, the agent was assigned a random initial position, orientation, and phase in the period of the unsteady wake.

Inspired by Monte Carlo approaches to random processes, we performed twenty independent training instances, each starting from a random policy (Table 2). Surprisingly, these twenty training instances consistently converged to two classes of policies, as evident from the evolution of the cumulative reward in Fig. 2B: nearly half the training instances converged to policies with reward shown in red and nearly half converged to policies with reward shown in blue (Table 2). In Fig. 2C, we show representative policies from each of the two classes of policies with colormaps representing the RL action 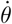 over the observation space (*s*_head_, *s*_tail_). The two classes of policies exhibit striking features: both are largely independent of the measurement at the head, and, although both converge during training, they exhibit opposite behaviors. The red policy, which we call hereafter *excitatory*, instructs the agent to turn in the same direction as the flow gradient at tail, that is, in the direction of increasing flow speed. The blue policy, which we denote *inhibitory*, instructs the agent to turn in the opposite direction to the flow gradient at tail, that is, in the direction of decreasing flow speed. Deploying either policy, the agent successfully tracks the wake to its generating source (Fig. 2D, Supp. Movie S.1). When applying the excitatory strategy, it ventures more into the wake, slaloming between vortices, similar to experimental observations in live animals [20, 24, 34, 42].

**Table 2:**
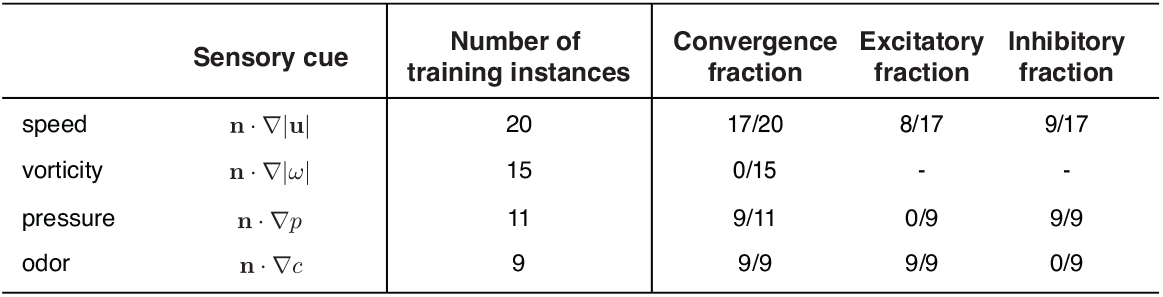
Training instances of RL policies based on different sensory cues. We conducted multiple training instances for four different signals transported by the flow field. The cumulative reward is shown in Fig. 2 and Fig. S1 based on representative training instances for each sensory cue.

**Figure 2:**
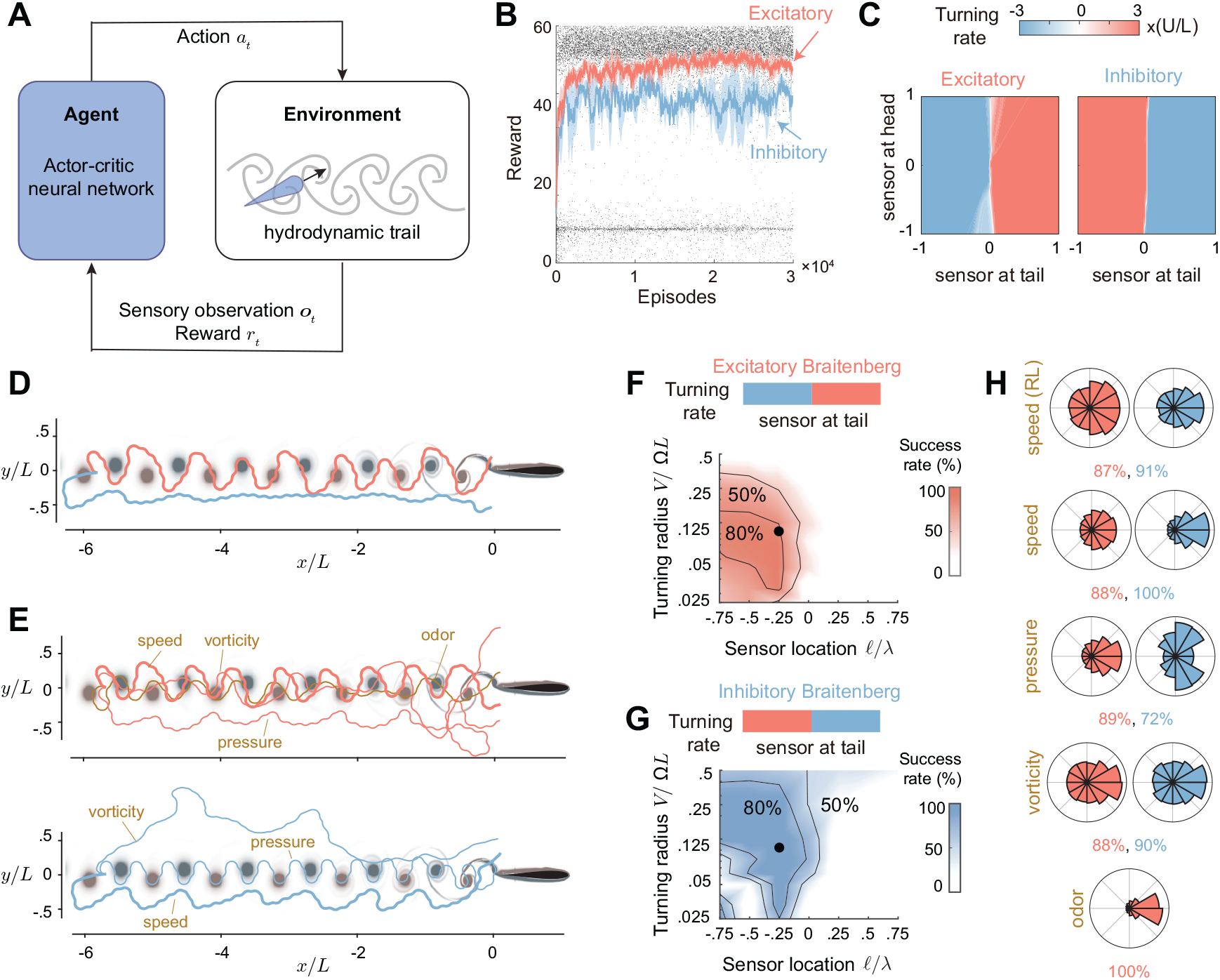
RL-policies and simpler Braitenberg-like policies for tracking hydrodynamic trails. **A**. To follow vortical trails, we train, using Deep RL, a swimmer that senses the flow gradients at its head *s*_head_ and tail *s*_tail_ (observations **o**_*t*_) and responds by modulating its angular velocity 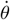 (action *a*_*t*_). **B**. Learning curves corresponding to twenty training instances, where nearly half converged to one of two classes of policies, respectively; moving average of the total reward is shown in red and blue; total reward for randomly chosen 10% of the training episodes is shown as black dots. Convergence rate of different training instances is reported in Table 2. **C**. Representatives of the two classes of converged policies shown as colormaps over the observation space (*s*_tail_, *s*_head_), normalized by the maximal value of the flow signal. Dominant features of these policies are captured by parsimonious Braitenberg-like strategies with excitatory and inhibitory responses to the flow signal measured only at tail (Eq. 2) and associated depiction at the top of panels F and G). **D**. Sample trajectories based on the excitatory (red) and inhibitory (blue) RL policies. **E**. Sample trajectories of the parsimonious representation of the excitatory (red) and inhibitory (blue) policies are superimposed based on different sensory cues. The same parameter values are used in RL-training 𝓁 = −0.25*L* and Ω = 3*U/L*. These trajectories are also compared in Supp. Movie S.3. Success rate as a function of sensor location 𝓁/*λ* and turning radius *V/*Ω*L* calculated over 76,500 test cases for each set of parameter values (𝓁/*λ, V/*Ω*L*) for **F**. excitatory and **G**. inhibitory policies in (2), with flow speed as sensory cue. Parametric study performed using the other sensory cues – pressure, vorticity, and odor – are reported in Fig. S2. **H**. Statistics on the gradient direction of the physical quantities measured by the onboard sensors along trajectories based on the RL or associated Braitenberg policies. The overall success rates over 76,500 test cases are listed for each sensory cue in red (excitatory) and blue (inhibitory).

### Learned policies are interpretable and analogous to Braitenberg’s Vehicles

These features of the RL policies – independence of *s*_head_ and nearly binary action (Fig. 2C) – suggest that they can be approximated analytically as bang-bang controllers [78] whose behavior depends only on the sensory measurement at tail *s*_tail_. Thus, without loss of significant properties, we discounted the sensory measurement at head *s*_head_ entirely and used 𝓁 and *s* to refer to the location and measurement of a single sensor, with negative 𝓁 implying sensor location at tail. These simplifications lead to analytic expressions of the two classes of RL policies,

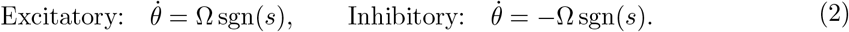

When implementing these simplified policies, the agent exhibited similar behavior to that dictated by the full RL policies (Fig. 2D,E, Supp. Movie S.1), while being readily interpretable and allowing for a rigorous stability analysis. These policies evoke Braitenberg’s vehicles, originally proposed as thought experiments for constructing complex behavior from simple sensorimotor responses to environmental stimuli [79]. Braitenberg’s vehicles have been applied to model animals’ tropotaxic behavior [80], including rheotaxis [81], phototaxis [82], chemotaxis [83]. In the simplest vehicle, two motors that drive the right and left wheels of a vehicle are either directly or inversely connected to two sensors, and each motor turns more as the stimulus to which the sensor is tuned increases. As in our strategies, the vehicle’s behavior depends on the differential signal between the two sensors. At zero difference, the vehicle moves straight ahead. When the motors are connected to sensors on the same side, the vehicle is aversive to stimulus and *turns away*, and when connected to opposite sensors, the vehicle is responsive to stimulus and *turns toward*. Likewise, in our excitatory policy, the swimmer turns toward larger flow speeds, while in the inhibitory policy, it turns away. In uniform flows, the swimmer swims straight.

### Robustness To Sensory Limitations

To evaluate the performance of the RL policies and their simplified counterparts in (2), we systematically tested them in a domain defined by a Cartesian grid starting downstream of the source. At each grid point, we considered equally spaced initial orientation between 0 and 2*π* and phase within the pitching period. In total, we tested 76,500 initial conditions for each policy (Methods). The excitatory and inhibitory RL policies achieved success rates of 87% and 91%, respectively, while the simplified counterparts in (2) achieved success rates of 88% and 100%, respectively, all tested in the same wake used during training.

We then evaluated the distributions of the terminal lateral locations *y/L* from the source (Fig. 3A) and the associated arrival time *t*_*f*_ (Fig. 3B). The excitatory policy is more accurate in honing in on the source (Fig. 3A), while the inhibitory strategy locates the source faster (Fig. 3B) because the corresponding trajectories meander less in the wake (Fig. 2D,E). By the same token, because it causes the agent to move inside the wake and slalom between vortices, the excitatory policy provides the agent with more opportunities to utilize the flow for efficient navigation (Methods, Fig. 3C,D). To assess the potential for efficient navigation, we calculated two metrics along each test trajectory. First, the flow agreement parameter 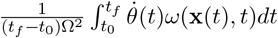 [84–86], measures the agreement between the agent’s angular velocity 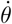 and the vorticity *ω*(**x**, *t*) of the ambient flow field at the agent’s location. Second, the thrust parameter 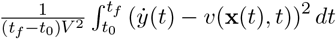 [86–89], evaluates how well the agent can utilize the transverse flow velocity *v*(**x**, *t*) to generate thrust. Compared to the inhibitory strategy, the trajectories instructed by the excitatory strategy have a more positive flow agreement and larger thrust parameters on average, (Fig. 3C,D). These trends are consistent between the RL policies and their simpler counterparts. Importantly, these findings show that by moving inside the wake, the excitatory strategy has more opportunities to harness the flow to swim more efficiently, consistently with empirical observations in fish [34].

**Figure 3:**
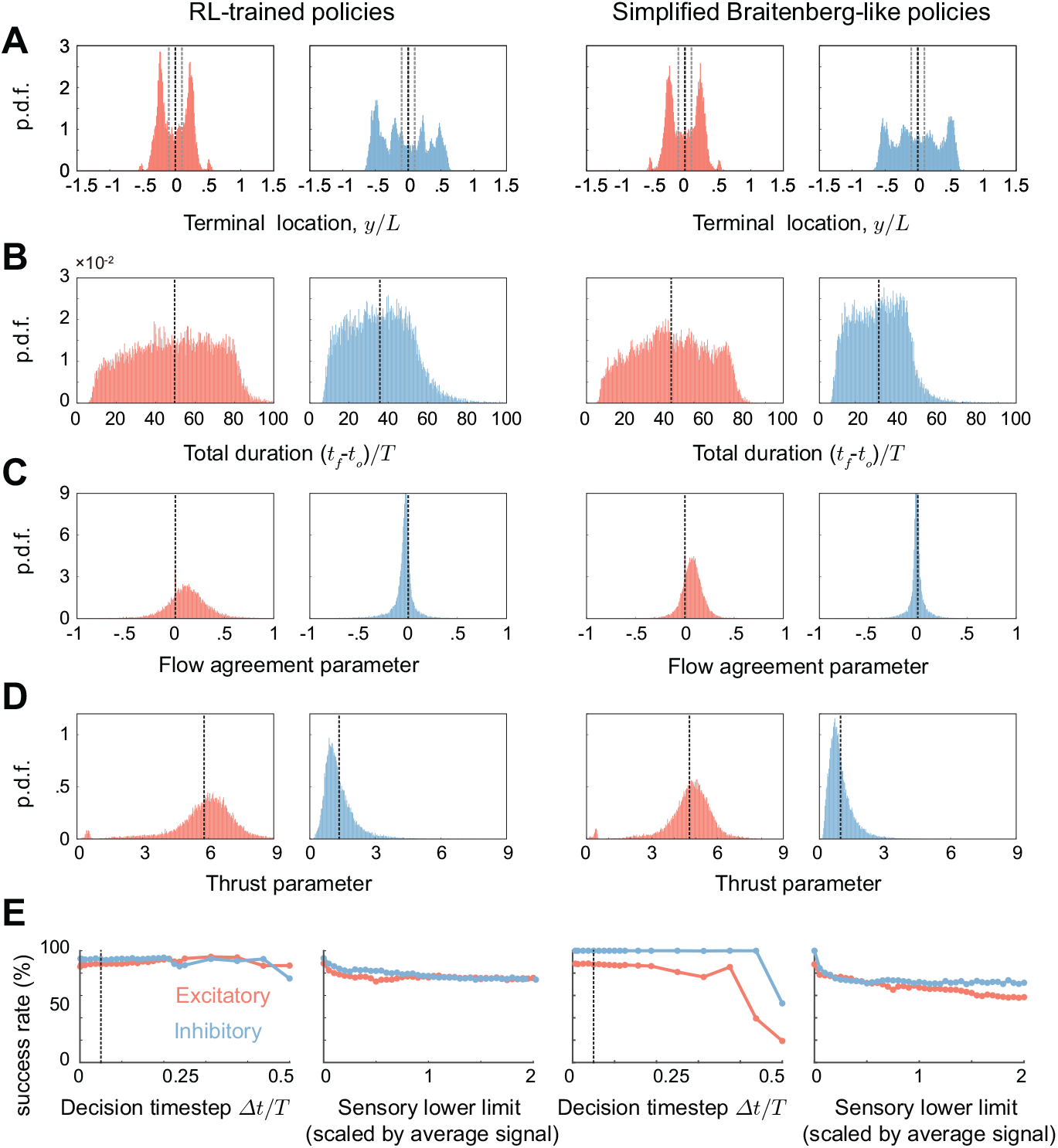
Performance of RL-trained policies and simpler counterparts. Each policy is tested on 76,500 test cases downstream of the source. Statistics are based on all trajectories, out of the 76,500 test cases, that succeed in moving upstream. The overall success rate for RL-trained policies and their simplified counterparts are 86.82% and 87.5%, respectively, for the excitatory policies (red), and 91.47% and 100% respectively, for the inhibitory policies (blue). **A**. Distribution of terminal location *y/L* at *t*_*f*_ defined as the time when *x/L* = 0. Dashed grey lines represent the tailbeat amplitude of the pitching foil. **B**. Distribution of total duration *t*_*f*_ − *t*_*o*_ to reach the source normalized by the period *T* = 1*/f* of the wake. The RL-trained policies are, on average, slower in reaching the terminal location than their simpler counterparts. **C**. Flow agreement parameter and **D**. thrust parameter along the successful trajectories. **E**. Success rate as a function of sensory limitations for the excitatory (red) and inhibitory (blue) policies: (leftmost) RL-trained policies are robust to increasing decision timestep Δ*t* up to 50% of pitching period *T*, that is, up to the agent changing direction only twice during a flapping period of the source. Dashed vertical line indicates the decision timestep used during training Δ*t* = 0.05*T*; (left middle) RL-trained policies are robust to sensory constraints where the agent cannot discern and respond to flow gradients below a lower limit. Success rate of the RL-trained policies decreases more gradually compared to their simpler counterparts (right middle and rightmost).

Lastly, to test the robustness of these policies under noisy and corrupt conditions, we challenged the agent in two major ways: (i) we decreased the rate at which it collects observations and responds to flow signals by increasing the decision timestep Δ*t*, and (ii) we imposed sensory limits *s*_min_ below which the agent is unable to sense the flow signal (Fig. 3E). Both the RL policies and the simpler versions in (2) exhibited remarkable robustness to these reduced abilities, with stronger robustness exhibited by the RL policies.

### Robustness to Agent Design

We next asked if the performance of the controller is sensitive to the specific agent design, including the choice of turning rate Ω and sensor placement 𝓁. During RL training, the agent’s turning rate was fixed at Ω = 3*U/L* and its sensor was placed at 𝓁 = −0.25*L*. Here, we set out to test the performance of each simplified policy in (2) over a broad range of sensor locations 𝓁/*L* ∈ [−0.75, 0.75] and turning rates Ω*L/U* ∈ [0.5, 10], corresponding to dimensionless turning radii *V/*Ω*L* ∈ [0.025, 0.5]. Using the same wake employed during training we conducted 76,500 tests for each combination of parameters (𝓁/*λ, V/*Ω*L*). The resulting success rates are shown in Fig. 2F and G, with the RL policy parameters indicated by black dots. Both the excitatory (red) and inhibitory (blue) policies achieved success rates of over 80% throughout nearly the entire parameter space, provided that sensory measurements were taken at the tail (*l <* 0). Importantly, because the performance of the RL-inspired strategies is insensitive to parameter values, an agent does not need to fine-tune its sensor location 𝓁 (provided *l <* 0) and turning radius *V/*Ω*L* to a specific wake type; thus, we expect the agent’s performance to be also robust to variations in the wake itself, a point that we revisit later.

### Robustness to Sensor Type

How do these trail-following strategies perform when different types of flow information are available to the agent? For example, if instead of measuring the difference in flow speed, the agent measures the difference in flow quantities such as pressure, vorticity, or the odor concentration carried by the flow field, how would that affect its ability to track hydrodynamic trails?

To investigate this question, we repeated the RL training using three distinct types of sensory cues: pressure, vorticity, and odor. For pressure sensors, the RL training converged to only inhibitory-like policies, while the odor sensors converged to only excitatory-like policies. Because the pressure field is on average weaker inside the wake, an inhibitory policy that instructs the agent to move to lower pressure is needed to stay inside the wake. Meanwhile, odor concentration is on average stronger inside the wake, thus, an excitatory policy keeps the agent in the wake. The RL training did not converge using vorticity measurements (Table 2 and Fig. S1). In the converged cases, the RL policies resemble bang-bang controllers similar to those proposed in (2).

In Fig. 2E, we show sample trajectories, starting from the same initial conditions, based on direct implementation of (2) using pressure, odor and vorticity sensors (see also Suppl. Movie S.3). Surprisingly, although vorticity measurements led to poor convergence in RL training, directly implementing this sensory cue in (2) allowed the agent to reorient and move upstream.

Importantly, when tested on the 76,500 sets of initial conditions in the wake of the source, we found that all four types of sensors – speed, pressure, vorticity, and odor sensors – exhibited high success rates in tracking the unsteady wake. This is surprising at first because when examining the distributions of gradient directions in the wake of the source (Fig. 1E), these distributions shows no bias towards the upstream direction. However, when computing the direction of the spatial gradient of the signal collected along each test trajectory (Fig. 2H), we found that for all sensors, the gradient directions exhibited a bias toward the upstream direction. This indicates that the motion of the agent shapes the signal it acquires.

We next probed the robustness of these results to variations in the agent’s design parameters (𝓁/*λ, V/*Ω*L*); Fig. S2 shows the success rates across the parameter space for each sensory cue. Both the excitatory and inhibitory strategies achieved high success rates of over 80%, over large ranges of the parameter space, provided that the sensory measurement is trailing the agent’s position (*l <* 0). Consistent with RL training, odor sensing functions only with the excitatory strategy. That is, it requires turning toward higher concentration. All other signals functioned well using either the excitatory or inhibitory policies, even the vorticity, for which the RL training did not converge. This is because the reward used in RL training requires the agent to accurately locate the source, whereas the success rate in Fig. S2 is based on the ability of the agent to move upstream, even if it ends at a lateral offset *y/L* from the source (trajectories in Fig. 2E). That is, vorticity measurements are viable to enable the agent to reorient and move upstream, as suggested in [90], but compromise the agent’s accuracy in locating the source. But even when tested based only on reorienting and moving upstream, the vorticity sensor underperformed compared to the other sensors – speed, odor, and pressure – in terms of range and robustness to agent design parameters and maximum success rate (Fig. S2). This, together with the failed training of the vorticity-sensitive agent, align with recent reports of the inefficacy of sensing vorticity in point-to-point underwater navigation [50, 51] and challenge approaches that advocate for sensing vorticity [90]. Sensing flow speed, pressure, or odor performs better in tracking flows. Because pressure sensors are most readily available commercially [31, 81, 91], the pressure-based strategy would be most amenable for immediate implementation in a robotic demonstration. Combining different types of sensors at once will be the topic of future studies and may lead to more robust trail-following behavior.

### Generalization to Diverse Hydrodynamic Trails

We next tested the performance of the simplified policies in (2) in wakes not seen during RL training. Specifically, we tested the policies in wakes generated by flow past pitching foils mimicking turning gait (Fig. 4A), intermittent swimming (Fig. 4D), and at different Strouhal and Reynolds numbers spanning St = 0.1 - 0.25 and Re = 500 - 5000 (Fig. 4C, Table 3). We also tested the policy in wakes generated by flow past a fixed cylinder, with Re = 200-500 (Fig. 4E, Table 3) and in a three-dimensional (3D) wake, adapted from [92], generated by an undulating body at Re = 2400 (Fig. 4B). For the 3D wake, we extended the simplified policies to control both yaw and pitch angles of the agent (Methods). In Table 3, we report a summary of the performance of each policy in each of these wakes. Because each wake has a different wavelength, we used the flow wavelength *λ* to scale the corresponding sensor location 𝓁 = −0.25*λ* in Fig. 1. Without any additional parameter tuning, the RL-inspired strategies achieved success rates of nearly 100% in guiding the agent to reorient upstream and track most of these hydrodynamic trails. In challenging scenarios, e.g. when direction of background flow is changing (Fig. 4A) or during intermittent flapping (Fig. 4D), in which the traveling wave structure is not always present in the entire time series, the excitatory strategy performed notably better than the inhibitory strategy by navigating inside the wake, where the traveling wave structure is more discernible. Thus, in addition to their parsimony and simplicity, the RL-inspired strategies are remarkably generalizable, allowing the agent to track a wide range of hydrodynamic trails, from thrust to drag jets to 3D wake behind undulatory, self-propelled swimmers (Fig. 4, Table 3, Supp. Movie S.2).

**Table 3:**
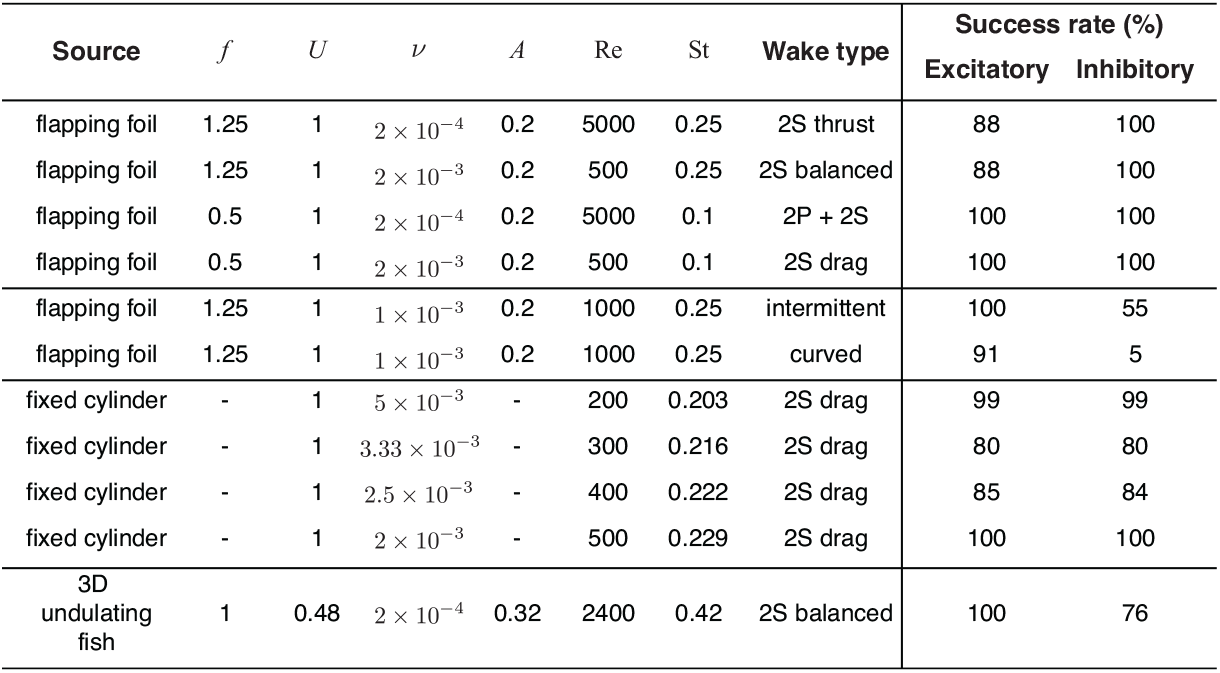
Success rate of excitatory and inhibitory strategies in CFD simulations of flows past stationary and moving bodies. Wake type depends on how many vortices are shed in a period and their spatial relationship [105, 106]. When two single vortices are shed in one period, such as Fig. 1C of the main text, the wake is called a 2S wake. If two pairs of vortices and two single vortices are shed during one period, such as in Fig. 4C of the main text, the wake is categorized as a 2P+2S wake. A von Kármán vortex street that generates a jet pointing upstream is a drag wake (Fig. 4E in the main text), while a reverse von Kármán vortex street is a thrust wake (Fig. 1C). The success rate of excitatory and inhibitory strategies of 𝓁 = −0.25*λ* and Ω = 3*U/L* are reported in the right two columns for different wake types.

| Source | $f$ | $U$ | $\nu$ | $A$ | Re | St | Wake type | Success rate (%) | |
| --- | --- | --- | --- | --- | --- | --- | --- | --- | --- |
|  |  |  |  |  |  |  |  | Excitatory | Inhibitory |
| flapping foil | 1.25 | 1 | $2 \times 10^{-4}$ | 0.2 | 5000 | 0.25 | 2S thrust | 88 | 100 |
| flapping foil | 1.25 | 1 | $2 \times 10^{-3}$ | 0.2 | 500 | 0.25 | 2S balanced | 88 | 100 |
| flapping foil | 0.5 | 1 | $2 \times 10^{-4}$ | 0.2 | 5000 | 0.1 | 2P + 2S | 100 | 100 |
| flapping foil | 0.5 | 1 | $2 \times 10^{-3}$ | 0.2 | 500 | 0.1 | 2S drag | 100 | 100 |
| flapping foil | 1.25 | 1 | $1 \times 10^{-3}$ | 0.2 | 1000 | 0.25 | intermittent | 100 | 55 |
| flapping foil | 1.25 | 1 | $1 \times 10^{-3}$ | 0.2 | 1000 | 0.25 | curved | 91 | 5 |
| fixed cylinder | - | 1 | $5 \times 10^{-3}$ | - | 200 | 0.203 | 2S drag | 99 | 99 |
| fixed cylinder | - | 1 | $3.33 \times 10^{-3}$ | - | 300 | 0.216 | 2S drag | 80 | 80 |
| fixed cylinder | - | 1 | $2.5 \times 10^{-3}$ | - | 400 | 0.222 | 2S drag | 85 | 84 |
| fixed cylinder | - | 1 | $2 \times 10^{-3}$ | - | 500 | 0.229 | 2S drag | 100 | 100 |
| 3D undulating fish | 1 | 0.48 | $2 \times 10^{-4}$ | 0.32 | 2400 | 0.42 | 2S balanced | 100 | 76 |

**Figure 4:**
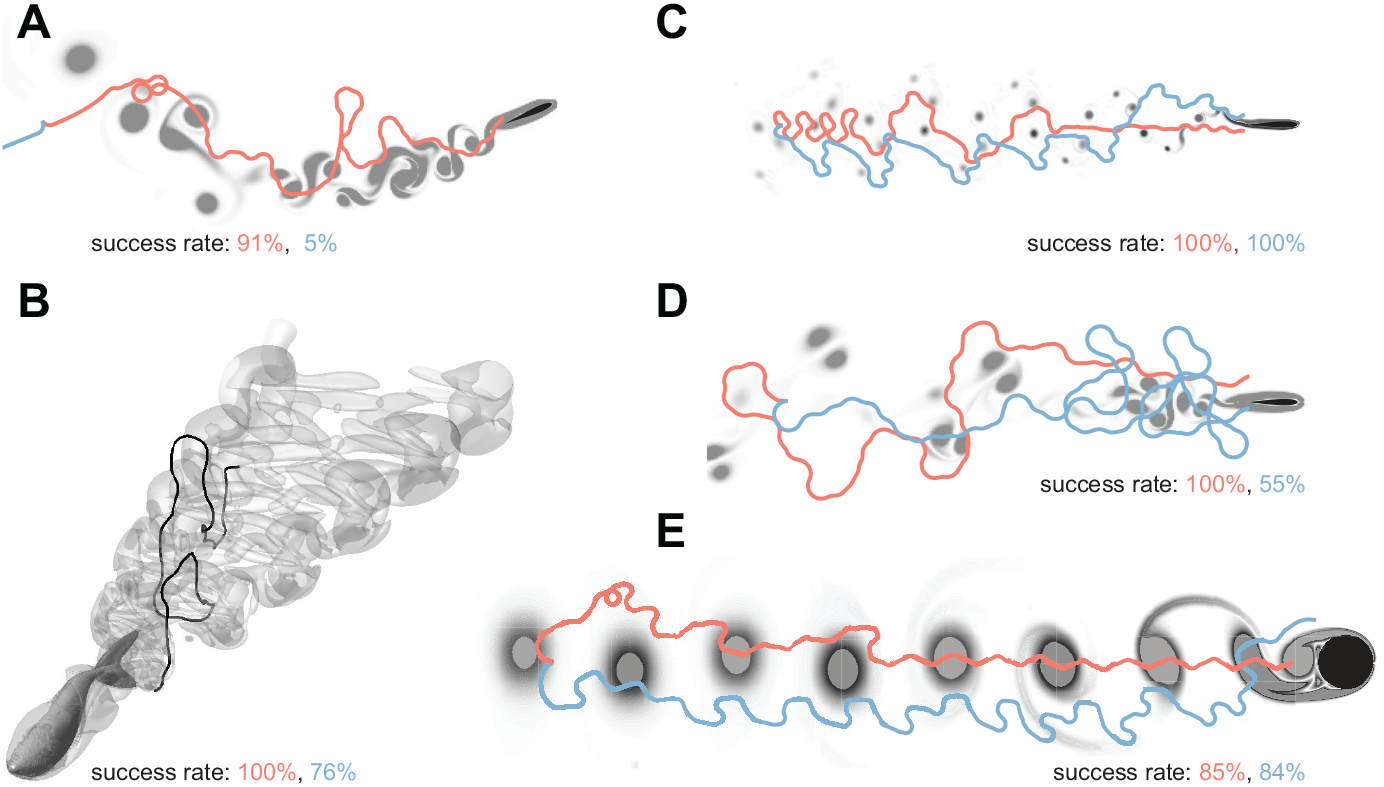
Generalization to unseen flows. Sample trail-following trajectories obtained by applying (2) to unseen wakes; agent successfully follows the wake of **A**. a pitching foil at Re = 1000 and St = 0.25 that changes direction of motion; **B**. 3D swimming fish [92]; **C**. pitching airfoil at Re = 5000 and St = 0.1; **D**. burst-and-cost pitching airfoil at Re = 1000, St = 0.25, and duty cycle 0.6; **E**. fixed cylinder at Re = 400. Success rates out of 76,500 test trajectories are listed for both the excitatory and inhibitory policies and reported in Table 3 for all hydrodynamic trails. Parameter values are set to Ω = ±3, and 𝓁 = −0.25*λ*. Sample movies are given in Supp. Movie S.2.

### Stability Analysis and Sensor Placement

The most salient feature among all tested wakes is the quasi-periodic and traveling-wave nature of the signal field (Fig. 1C). But how does this feature contribute to the success of the RL policies and their simpler Braitenberg-like analogs? More importantly, why is the sign of control gain not important, with inhibitory or excitatory strategies working equally well, while the sensor placement is critical? To address these questions, we analyzed the stability of the RL-inspired strategies in a 1D traveling-wave signal field of the form *g*(*x, t*) = *A* cos(2*π*(*x* + *λft*)*/λ*), which is a general solution of the wave equation *g*_*tt*_ = *λ*^2^*f* ^2^*g*_*xx*_. This 1D wave solution is the simplest representation of the flow signal left by a fish-like swimmer [86, 89].

The excitatory and inhibitory strategies are related by a change of sign from Ω to −Ω, which, in the traveling wave signal field, is equivalent to a translation in either space *x* → *x*+*λ/*2 or time *t* → *t* + 1*/*(2*f*). It thus suffices to analyze, say, the excitatory strategy with the understanding that the asymptotic behavior of both strategies is similar.

Trail tracking in this 1D traveling wave signal field amounts to moving upstream in the direction of *θ* = 0, opposite to the direction of wave propagation. Numerical simulations show that upstream motion in this signal field depends on the sensor location 𝓁 (Fig. 5A). To analyze the problem analytically, we introduced a coordinate transformation *z* = *x* + *λft*, where *z* represents a Lagrangian point moving with the traveling wave, and substituted *z* and *g*(*z*) = *g*(*x* + *λft*) into (1) and (2) to eliminate explicit dependence on time,

**Figure 5:**
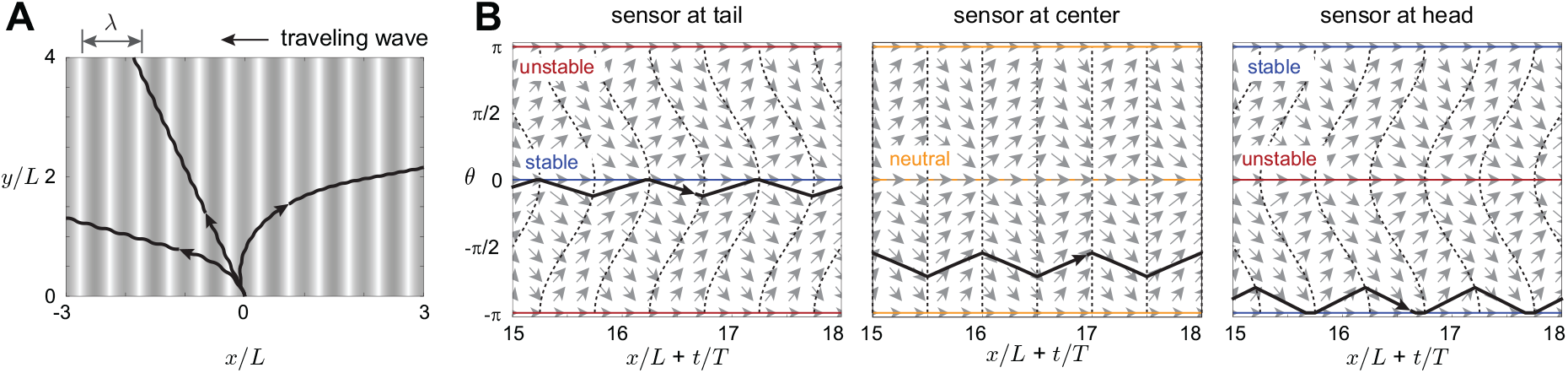
Stability analysis. **A**. Sample trajectories in 1D traveling signal field *A* cos(2*πx/λ* + 2*πft*) by three agents with sensor placements at tail (𝓁 = −0.25), center (𝓁 = 0), and head (𝓁 = 0.25), all starting from the same initial condition (*x, θ, t*)|_0_ = (0, 0.55*π*, 0). **B**. Phase portraits on the phase space (*x* + *λft, θ*) for sensor at tail (𝓁 = −0.25), center (𝓁 = 0), and head (𝓁 = 0.25). Trajectories are superimposed after reaching the quasi-steady state. Parameter values are set to swimming speed *V* = 0.25 and turning rate Ω = 1.

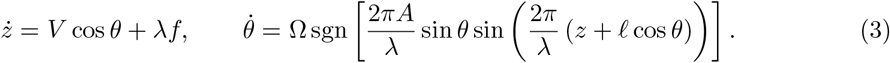

The system in (3) has two moving equilibria at *θ* = 0 (moving upstream) and *θ* = ±*π* (moving downstream). The stability of these equilibria depends on the sensor location. Fig. 5B depicts the phase portraits of (3) over the phase space (*z, θ*). Dashed black lines indicate level sets where 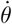 is 0, with phase velocity 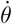 alternating directions between adjacent level sets. The linear stability at these equilibria *θ* = 0, ±*π* varies periodically with *z*. The shape of the level sets ensures that when *l <* 0, trajectories travel longer distance in the regions where the phase velocity points toward *θ* = 0; the upstream equilibrium is thus stable. For *l >* 0, trajectories travel longer in the regions where the phase velocity points toward *θ* = ±*π*; the downstream equilibrium is stable. At the bifurcation point 𝓁 = 0, both equilibria are neutrally stable.

This finding, that trail following is stable when sensors are placed at the tail, is intriguing given that biological evidence indicates that flow sensors are mostly located at the head, such as in the whiskers of harbor seals [24] and in the lateral line system in fish where flow-sensitive neuromasts are denser at the head [74, 75]. Indeed, physics-based [74, 75] and data-driven [93, 94] models suggest that the flow signal measured at the head is more informative in decoding flow information compared to the flow signal measured at the tail. However, recent behavioral experiments found that disabling the posterior portion of the lateral line is harmful to certain behaviors, such as fish rheotaxis [90, 95]. Our study offers a potential hydrodynamic explanation for why a sensor at the tail is crucial for hydrodynamic tracking and rheotaxis. Another perspective is that a trailing sensor is mathematically equivalent to a sensor at the head with a time delay *τ* between sensing and response. Indeed, in aquatic animals, there is a time delay in the neural system between sensory input and behavioral response [96]. To explore this hypothesis, we collected biological data (Table S1) and mapped their time delay *τ* to sensor placement 𝓁 ≈ *V τ*. Remarkably, organisms that inhabit and track hydrodynamic trails at comparable Reynolds numbers [20, 21, 24, 42] lie within a parameter range compatible with our model predictions for stable and robust trail following (Fig. S5).

### Sensing Longitudinal Gradients

Our trail-tracking policies relied on sensing the transverse gradient of the signal field **n** · ∇. Do these policies work when sensing the longitudinal gradient of the signal field **t** · ∇? In Fig. 6A, we show in simulations that measuring the longitudinal gradient of the flow speed allows effective trail tracking in vortical flows. Through a rigorous stability analysis similar to that presented above, we found that sensing the longitudinal gradient of the flow signal results in stable upstream motion when the sensor is placed at the tail (Fig. S4). When evaluating other sensory cues such as odor, pressure, and vorticity, we found that their performance was suboptimal compared to flow speed (Fig. S3). We next placed the longitudinal sensor at the center 𝓁 = 0 and allowed a time delay *τ* between sensing and response (Fig. 6B). Again, we tested this sensory strategy across all 76,500 initial conditions; the overall success rate of the corresponding policy was nearly as good as that with the sensor placed at the tail.

**Figure 6:**
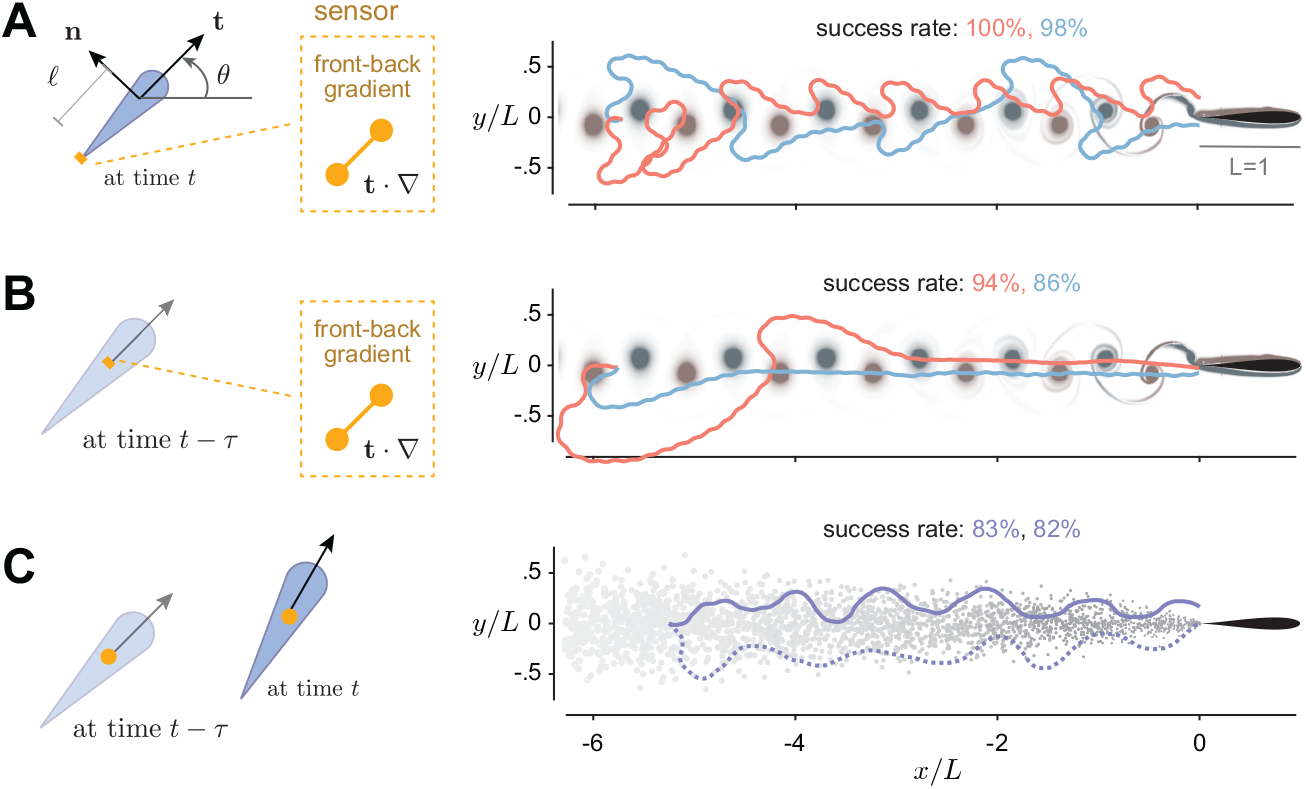
Simple and versatile strategies for tracking vortical and turbulent wakes. Sample trajectories and success rates of simple yet adaptable trail-following strategies: **A**. an agent measuring longitudinal gradient of flow speed at tail, with 𝓁 = −0.25 and Ω = 3, is equally successful in tracking vortical wakes as the agent measuring lateral gradient in Fig. 2. **B**. an agent, with sensor collocated at the agent’s position, measuring longitudinal gradient at a time delay *τ* = 0.1, is also equally successful in tracking vortical wakes. **C**. An agent measuring temporal difference in the signal itself, not the signal gradient, is successful in tracking a turbulent wake. Here, *τ* = 1 and Ω = 2. The trajectories show the classic “cast and surge” behavior [57, 58]. As a baseline for comparison, the success rate in [19] of RNN agent tracking signal field of constant direction is about 75%. Movie of the trajectory is shown in Supp. Movie S.2.

### Bridging Trail-Tracking Strategies from Vortical to Turbulent Trails

To extend these policies to turbulent plumes where measuring spatial gradients of the signal field is not viable [18, 19], we considered an agent that measures the odor signal itself at time *t s*(*t*) = *g*(**x**(*t*), *t*) and compares it to that measured at a previous time *t* − *τ s*(*t* − *τ*) = *g*(**x**(*t* − *τ*), *t* − *τ*). Following one strategy, the agent rotates with angular velocity Ω when *s*(*t*) *> s*(*t* − *τ*) and −Ω when *s*(*t*) *< s*(*t*−*τ*); and the other strategy does the opposite (9). In Fig. 6C, we show that both strategies achieve similar performance with over 80% success in tracking the turbulent plume. The success of these simple strategies in a turbulent plume stems from the fact that, despite the absence of a traveling wave structure with well-defined frequency and/or wavelength, odor patches are consistently advected downstream by the mean flow (Fig. 1B, [12]).

## Summary and Implications to Underwater Robotics

Using deep reinforcement learning, we discovered two policies for tracking biologically relevant hydrodynamic trails at intermediate Reynolds numbers. The policies rely only on local and instantaneous flow sensing, and can be expressed in terms of two parsimonious, interpretable, and generalizable control strategies, where the swimmer measures locally a differential flow signal and responds by turning either toward or away from the direction of stronger signal (Fig. 2). These remarkably simple strategies depend only on two dimensionless parameters – the minimum turning radius *R* = *V/*Ω*L* of the agent and the sensor location 𝓁/*L*. Through rigorous stability analyses, we proved that these strategies are stable in signal fields with traveling wave character, provided posterior sensor placement 𝓁/*L <* 0 (Fig. 5), and we offered both biological and hydrodynamic explanations for these findings. Using Monte Carlo simulations, we demonstrated that these strategies carry over to unfamiliar wakes (Fig. 4, Table 3) and to sensors that probe different types of flow signals (Fig. 2D,E and Fig. S2). Moreover, inspired by these strategies, we hand-crafted a simple strategy to track turbulent plumes, where the spatial gradient of the signal is not available, by responding to the temporal difference of the signal field (Fig. 6B).

Our strategies outperform existing sensory and response mechanisms for trail tracking (Table 1) in three assessment metrics – interpretability, versatility, and adaptability to different flow currents. Importantly, our strategies are remarkable for their minimal sensory requirements. Sensing is expensive! Sensory limitations are thought to be the greatest challenge preventing robotic systems from achieving performance on par with their animal counterparts [97]. Our findings show that trail tracking of flow currents can be robustly achieved with minimal requirements on sensory abilities and memory.

Our work offers a powerful framework for designing robust and adaptable underwater navigation systems. Case in point, we employed a three-link fish model [98, 99] and developed, based on the RL-inspired excitatory and inhibitory strategies, controllers that directly mapped local flow signals to body deformations, much like Braitenberg’s vehicles that linked signal intensity to wheel rotation [79]. Numerical tests confirmed the success of the three-link fish in tracking a traveling wave signal (Suppl. Movie S.1). Further analysis and experimental validation of these predictions will be the topic of future research [29, 31, 32].

Our findings also guide the strategic placement of sensors in underwater robotic systems. Surveying existing systems [36, 100–102] (Table S1), we found that robots capable of large ranges of body deformations exhibited turning radii comparable to those of swimming organisms [36, 100, 101] (Table S1). Then, using our results (Fig. 2F and G) that quantify the success in trail following over the space of sensor location and turning radius, we predicted ranges of sensor locations (Fig S5) for which these robotic systems would exhibit success rates exceeding 95%. These predictions provide valuable tools for designing and optimizing the sensory layout of future underwater robotic systems and testing them in real-world scenarios, thus advancing the development of underwater technologies [29, 31, 32].

## Methods

### Simulations of Flow Environment

#### Laminar Plume

An odor-emitting source with continuous emission rate *R* is located at point **x**^*^. Time evolution of the odor concentration field is composed of advection of background flow **u**(**x**, *t*) and diffusion at constant diffusive constant *D*

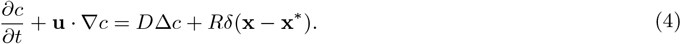

In laminar plume, the flow is a uniform background flow **u**(**x**, *t*) = *U* = 1. Thus, Péclect number is defined as the ratio between the advection rate and diffusion rate *Pe* = *LU/D*. The solution of (4) is given by

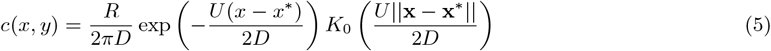

where *K*_0_ is the modified Bessel function of the second kind of order zero [18]. The parameters are chosen as: odor emission rate *R* = 0.25, uniform background flow velocity *U* = 1, and diffusion rate *D* = 0.1. These result in a Péclect number Pe = 10.

#### Turbulent Plume

We employed a particle-based two-dimensional plume model [19, 66]. The plume model simulates the random release of odor packets from a source with a fixed initial radius *r*_0_ and initial strength *c*_0_. A turbulent wind transported the odor packets, and spread out in all directions due to radial diffusion. The concentration decreases with the increases in packet size

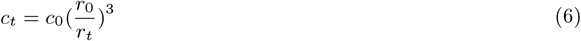

The turbulent wind is simplified to the combination of a uniform background flow of velocity *U* = 1 and random perturbations with strength *σ*. These parameters are chosen as emission rate *R* = 1, initial radius and concentration of the plume *r*_0_ = 0.01 and *c*_0_ = 1, random perturbations cross-wind *σ* = 0.005, and diffusion rate 0.001.

#### Vortical Trail

Hydrodynamic trails were generated from computational fluid dynamics (CFD) simulations and used in this study (Table 3). These simulations included two-dimensional (2D) wakes generated by flows past an oscillating airfoil, 2D wakes generated by flows past a fixed cylinder, and a three-dimensional (3D) wake generated by flow past an undulating fishlike body.

In the CFD simulations, the spatial-temporal evolution of the flow field (velocity **u**(**x**, *t*), pressure *p*(**x**, *t*), and vorticity *ω*(**x**, *t*) = ∇ × **u**) is governed by the incompressible Navier-Stokes equations, given in dimensionless form as

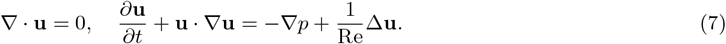

The boundary conditions are uniform velocity inlet and no-slip boundary conditions on the surface of the structure. The flow fields are solved using immersed boundary method (IBM) to deal with the moving boundary [68, 69, 107–109].

The 2D simulations were conducted using an open-source fluid solver, IBAMR [67–69]. The airfoil is a symmetric Joukowsky airfoil of chord length *L* and maximum thickness 0.12*L*, undergoing pitch oscillations *α*(*t*) = *α*_0_ sin(2*πft*) about a pivot point located at 0.25*L* from its leading edge, where *α*_0_ is the amplitude and *f* is the pitch frequency. The peak-to-peak tailbeat amplitude is *A* = 1.5*f* sin *α*_0_. The parameter values are summarized in Table 3.

We chose a rectangular computational domain of size [−24, 8] × [−8, 8], and placed the pivot point of the airfoil at the origin (0, 0) of the domain. The coarsest Eulerian mesh is a uniform 128 × 64 Cartesian grid. There are 3 layers of refinement mesh, with refinement ratio = 4 for each layer. The refinement criterion is based on the vorticity of the wake. The simulation timestep is Δ*t* = 1 × 10^−4^. The types of simulated wakes are listed in Table 3. The 3D wake generated by flow past the undulating body is based on ViCar3D [92, 108, 110]. Details of the computational setup are reported in [92].

To consider the concentration field advected by the flow field, we placed an odor-emitting source at the trailing edge of the airfoil and solved the advection-diffusion equation for the concentration field advected by the flow field (4). The Péclect number in Fig. 1C is 10, 000.

### Reinforcement Learning

#### Training the Agent using Deep Reinforcement Learning

We implemented the clipped-advantage proximal policy optimization (PPO) method proposed by [76] for our RL training as in [98]. PPO maximizes a surrogate objective that clips off unwanted changes when the policy deviates too much from the policy of the previous experience to ensure faster and more robust convergence. Two neural networks, actor and critic (both 64 × 32), which approximate the mean of the policy and value function of the state, respectively, are being updated during the training process. We refer readers to the original reference cited above as well as the OpenAI’s documentation of the PPO algorithm [111], their baseline implementations [112] and previous paper from our group [98] for a thorough explanation of the theory and details behind this method. In this work, both actor and critic are represented and fitted by 64 × 32 Deep Neural Networks (DNNs).

We trained the agent to track the hydrodynamic trail from repeated experiences interacting with the flow environment. The RL agent took as observations *o* = (*s*_head_, *s*_tail_) two sensory measurements taken at head and tail, and provided as action a noisy turning rate 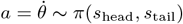.

To limit the action space, we imposed a threshold value Ω on the maximal angular velocity 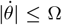. This, in turn, constrained the minimum turning radius 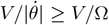 of the agent. The decision timestep Δ*t* is chosen as 0.05*T*, where *T* is the period of pitching. When it reached the source in the streamwise direction, the agent received a sparse reward of 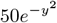, decaying with the lateral distance *y* from the source. Additionally, to encourage initial learning, the agent received a dense shaping reward at each decision timestep Δ*t*, −*d*(*t*) + *d*(*t* −Δ*t*), smaller in magnitude, that depended on its relative change in distance *d*(*t*) to the trailing edge of the source generating the flow, yielding a reward 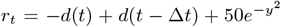. The reinforcement learning algorithm is trying to maximize total discounted return, 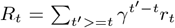. Thus, the algorithm tries to find a time optimal policy by itself.

#### Evaluating Policy Success Rate

To test the performance of the strategies, and to standardize these tests across different wakes and sensory cues, we designed a test domain consisting of a rectangular grid starting at a distance *L* downstream from the trailing edge of the airfoil and spanning a 9*L* × 3.2*L* range in the *x* and *y* directions, with grid size 0.2*L*. This grid provides 45 × 17 = 765 distinct initial positions (*x*(0), *y*(0)) of the agent. At each grid point, we explored 20 equally spaced initial orientations *θ*(0) between [0, 2*π*) and 5 equally spaced initial phases in a pitching period [0, 1*/f*). That is, for each wake and each policy, we run 76, 500 tests.

In the three-dimensional flow field (Fig. 4B), the initial condition is composed of 6 independent variables (*x*(0), *y*(0), *z*(0), *θ*(0), *ψ*(0), *t*(0)). Doing a grid search is infeasible under this scenario, so we generated 90, 000 random initial conditions to evaluate the performance.

### Handcrafted Strategies for Hydrodynamic Trail Following

#### Simplified Strategies based on Local Sensing of Signal Gradient

We devised Braitenberg-like strategies where the agent simply responds to the local gradient of the signal field *g*(**x**, *t*). The gradient can be taken in either the normal or tangential direction, namely

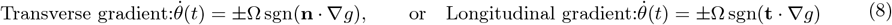

An excitatory strategy responding to the transverse gradient of concentration is similar to a gradient ascent; it works perfectly in laminar plumes. In turbulent plumes, measuring spatial gradients is not feasible given the sporadic and random distribution of the signal. In vortical wake, multiple local minima and maxima exist in the concentration field, and thus measuring normal and longitudinal gradients both work as long as the sensor is not collocated with the agent’s position, but placed at the tail, behind the agent.

#### Strategies Based on Temporal Difference of Detected Signal

When measuring spatial gradient is not applicable, the sensory control strategies are generalized by substituting spatial gradient with finite temporal difference. Namely, at each timestep *t*, it compares the current signal *g*(**x**(*t*), *t*) with a previous the signal at previous time *g*(**x**(*t* − *τ*), *t* − *τ*) and turns based on whether it is bigger or smaller than the current signal:

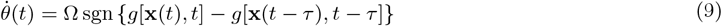

The temporal difference of signal is an approximation of the longitudinal gradient of the signal field plus a time delay. Although the strategy does not work in the laminar plume and vortical wake, it achieves a high success rate in the turbulent plume as illustrated in Fig. 6C. In turbulent plume, Ω *>* 0 and Ω *<* 0 generate similar success rates (Fig. 6), but do not intuitively map to excitatory or inhibitory strategy since they are responding to temporal difference instead of lateral gradient.

#### Application of Braitenberg Strategies in 3D wakes

In 3d, the agent’s position vector **x** is expressed in terms of the coordinates **x** = (*x, y, z*) in the fixed inertial frame (**e**_*x*_, **e**_*y*_, **e**_*z*_). We affixed a body frame (**t, n, b**) to the agent, where **t** is a tangential unit vector in the agent’s swimming direction and **n** and **b** are the normal and binormal unit vectors. We used Euler angles (*θ, ψ, ϕ*) to describe the orientation of the agent. We excluded spinning around **t** by setting *ϕ* = 0, and controlled the agent’s heading angle *θ* and *ψ* based on the gradient of the signal field in the **b** and **n** directions, respectively. Thus, in lab frame (**e**_*x*_, **e**_*y*_, **e**_*z*_), the body frame is expressed as **t** = (cos *ψ* cos *θ*, sin *ψ* cos *θ*, − sin *θ*), **n** = (− sin *ψ*, cos *ψ*, 0), **b** = (cos *ψ* sin *θ*, sin *ψ* sin *θ*, cos *θ*). In this c, the agent’s equations of motion are written as

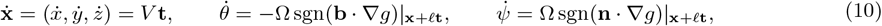

where *g*(**x**, *t*) is the signal field. In the case shown in Fig. 4B, the sensory cue is chosen to be the flow speed |**u**|.

### Biological Data

We sought to evaluate our predictions of the success rate of the RL-inspired policies over the parameter space of turning radius *V/*Ω*L* and sensor location 𝓁/*L* expressed in terms of time delay *τ* between sense and response, 𝓁/*L* ≈ *V τ/L*, given parameters achievable by biological and robotic systems. We thus gathered data on the turning radius and swimming speeds of aquatic organisms and underwater robotic vehicles. Fish data are based on the review paper by Domenici *et al*. 1997 [113] and other sources [114–125]. Data for sea lions, which are close relatives of harbor seals, came from [126].

For the time delay *τ* between mechanosensing and motor response of aquatic animals, we relied on measurements of the latency of fish evasion response to a mechanical stimulus [127]. The time delay between stimulus and body flexion, indicating delay in neural activity, ranged from 5 to 150 ms. Similarly, in [128], the authors measured the stimulus-to-response time exhibited by hatchling *Xenopus* tadpoles and found the response time is about 70 ms. For harbor seals, [129] measured the reaction time of harbor seals to acoustic stimulus, and found the reaction time of body motion ranged from 188 – 982 ms.

Some of these data were available in the min-max range (e.g. [127, 129]), which we converted to mean and standard deviation using the range rule std = (max − min)*/*4 [130]. The complete dataset is available in the Table. S1.

**Table S1:** Survey of relevant biological and robotic data from existing literature.

| Biological Systems` |  |  |  |  |  |  |
| --- | --- | --- | --- | --- | --- | --- |
|  | Common name | Body length (m) | Swimming speed (m/s) | Turning radius (m) | Reynolds number | Reference |
| Fish | Pacific trout | 0.33 | 0.165 | 0.13 | $5.4 \times 10^5$ | Weihs 1973 |
| | Dolphinfish | $2 \pm 0.3$ | $1.5 \pm 0.5$ | $0.33 \pm 0.08$ | $3.0 \times 10^6$ | Domenici & Blake 1997 |
| | Knifefish | 0.113 | $0.95 \pm 0.17$ | 0.0062 | $1.1 \times 10^6$ | Domenici & Blake 1997 |
| | Anglefish | 0.073 | $0.79 \pm 0.068$ | $(4.7 \pm 0.15) \times 10^{-3}$ | $5.7 \times 10^5$ | Domenici & Blake 1997 |
| | Pike | $0.51 \pm 0.044$ | $0.221 \pm 0.035$ | $0.046 \pm 0.004$ | $1.1 \times 10^6$ | Domenici & Blake 1997 |
| | Smallmouth bass | $0.31 \pm 0.07$ | $0.8 \pm 0.2$ | $0.034 \pm 0.014$ | $2.5 \times 10^6$ | Domenici & Blake 1997 |
| | Rainbow trout | $0.12 \pm 0.0035$ | $0.35 \pm 0.01$ | $0.021 \pm 0.003$ | $4.1 \times 10^5$ | Domenici & Blake 1997 |
| | Yellowtail | 0.19 | $0.77 \pm 0.094$ | 0.043 | $1.5 \times 10^6$ | Domenici & Blake 1997 |
| | Yellowfin tuna | $0.85 \pm 0.048$ | $0.68 \pm 0.11$ | $0.40 \pm 0.17$ | $5.7 \times 10^5$ | Domenici & Blake 1997 |
| Pinniped | Sea Lion | $1.8 \pm 0.085$ | $2.89 \pm 0.77$ | $0.41 \pm 0.14$ | $5.2 \times 10^7$ | Fish et al. 2003 |
| Underwater robotic systems |  |  |  |  |  |  |
| Underwater robots | Multi link fish | 0.45 | 0.25 | 0.09 | $1.1 \times 10^6$ | Li et al. 2020 |
| | Multi link fish | 0.34 | 0.063 | 0.1 | $2.1 \times 10^5$ | Hirata et al. 2000 |
| | Multi link fish | 0.65 | 0.19 | 0.22 | $1.2 \times 10^6$ | Yu et al. 2008 |
| | Driven by caudal fin | 1.23 | 1.1 | 1.8 | $1.3 \times 10^7$ | Liang et al. 2011 |
| | Driven by propeller | 1.6 | 1.1 | 4.2 | $1.8 \times 10^7$ | Liang et al. 2011 |
| Response time in biological systems |  |  |  |  |  |  |
|  | Response time (ms) |  | Reference |  |  |  |
| Fish | 5 - 150 |  | Domenic & Hale 2019; Roberts et al. 2019 |  |  |  |
| Seal | 188 - 982 |  | Kastelein et al. 2019 |  |  |  |

**Figure S1:**
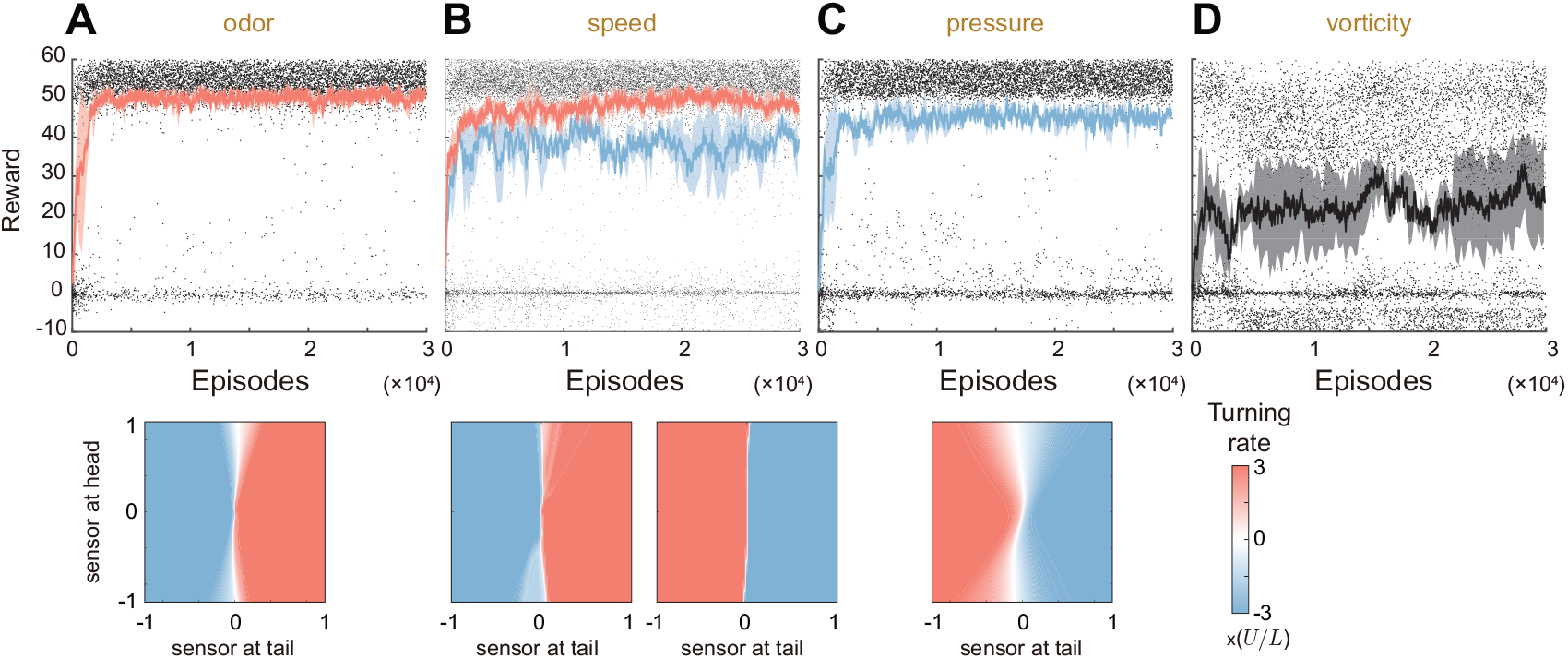
RL-training for different sensory cues. **A**. Lateral gradient of odor concentration. **B**. Lateral gradient of flow speed. **C**. Lateral gradient of pressure. **D**. Lateral gradient of vorticity magnitude. The convergence rates over multiple training instances are reported in Table 2. In **A** and **C**, RL training converges to excitatory and inhibitory strategies, respectively. In **B**, different training instances converge to either excitatory or inhibitory strategies. In **D**, the training does not converge. The insets show the converged RL policy by plotting the action 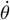 as a colormap over the observation space (*s*_tail_, *s*_head_). These colormaps exhibit similar features, though not as pronounced, to those obtained in Fig 2C of the main text: to first order-approximation, the action 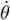 is mostly independent of the signal *s*_head_. We thus approximate these strategies following (2). Parameter values: 𝓁_tail_ = −0.25*L, l*_head_ = 0.25*L*, Ω = 3*U/L*. The flow field used during training is the same as that in Fig. 2 of the main text.

**Figure S2:**
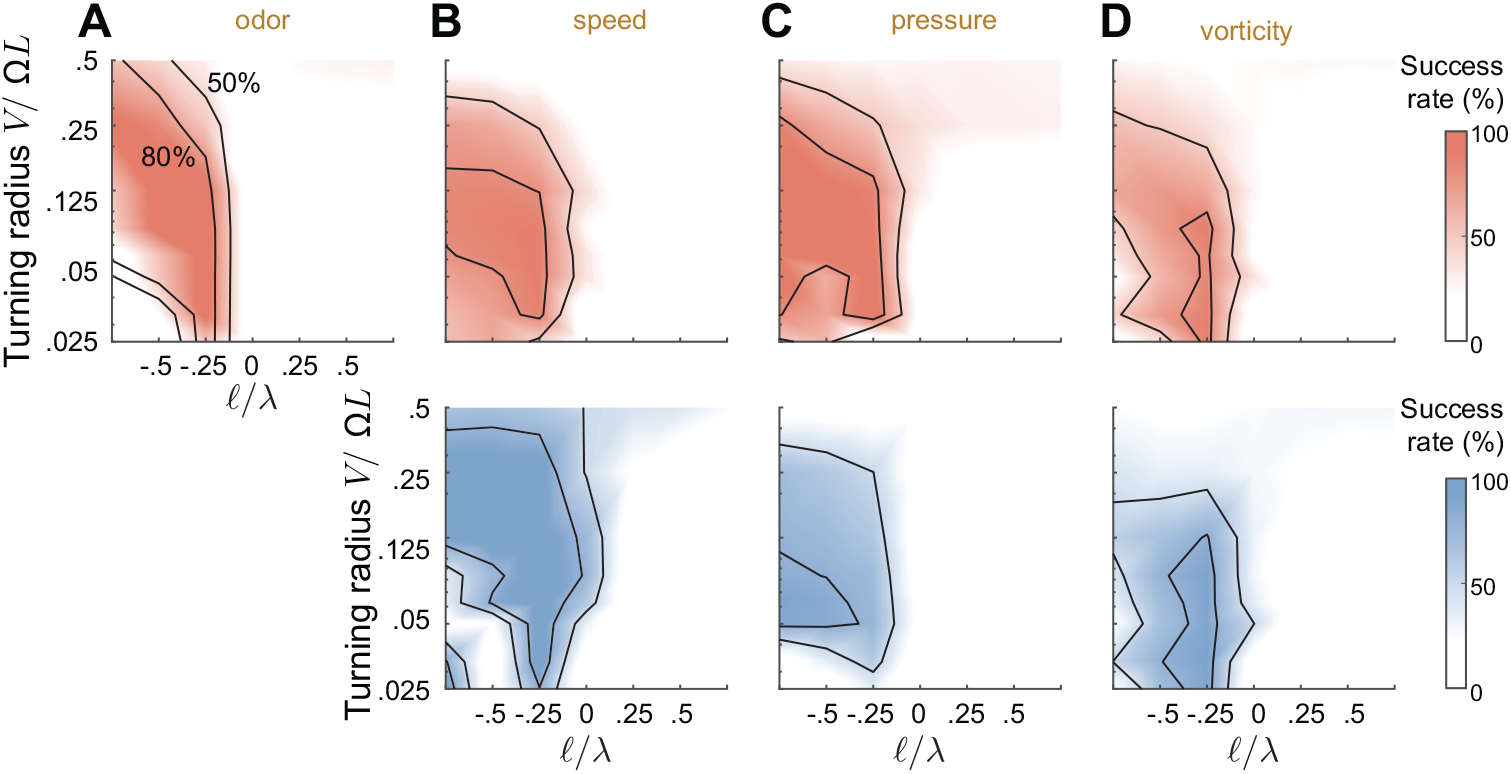
Parametric study for different sensory cues. Success rate of the agent as a function of sensor location 𝓁/*λ* and turning radius *R* = *V/*Ω*L* for different sensory cues with excitatory (red) and inhibitory (blue) RL-inspired strategies. Four different sensory cues are tested: **A**. odor, **B**. flow speed, **C**. pressure, and **D**. vorticity.

**Figure S3:**
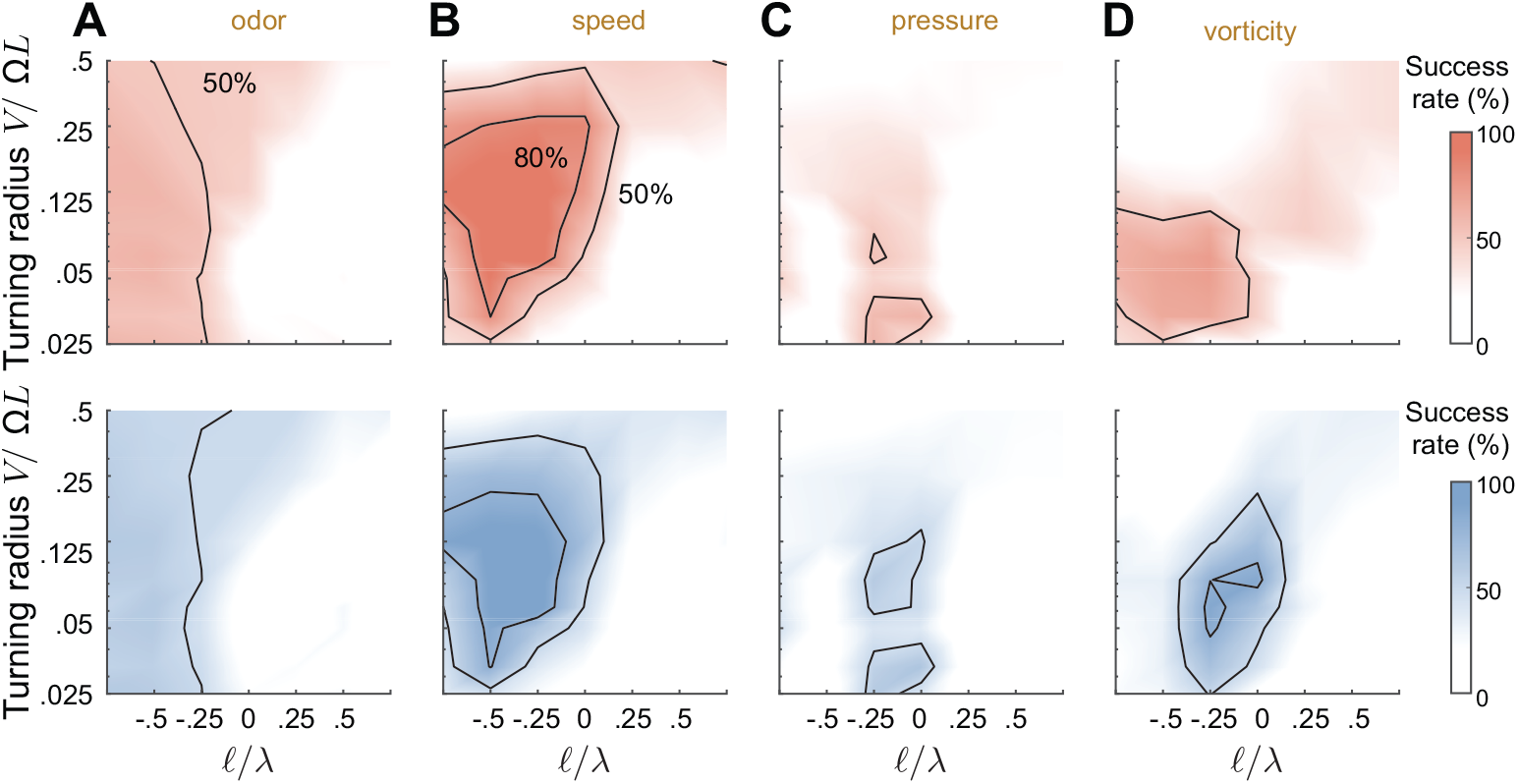
Parametric study for different sensory cues using gradient in longitudinal direction. Success rate of the agent as a function of sensor location 𝓁/*λ* and turning radius *R* = *V/*Ω*L* for different sensory cues with excitatory (red) and inhibitory (blue) RL-inspired strategies. Four different sensory cues are tested: o**A**. odor, **B**. flow speed, **C**. pressure, and **D**. vorticity.

**Figure S4:**
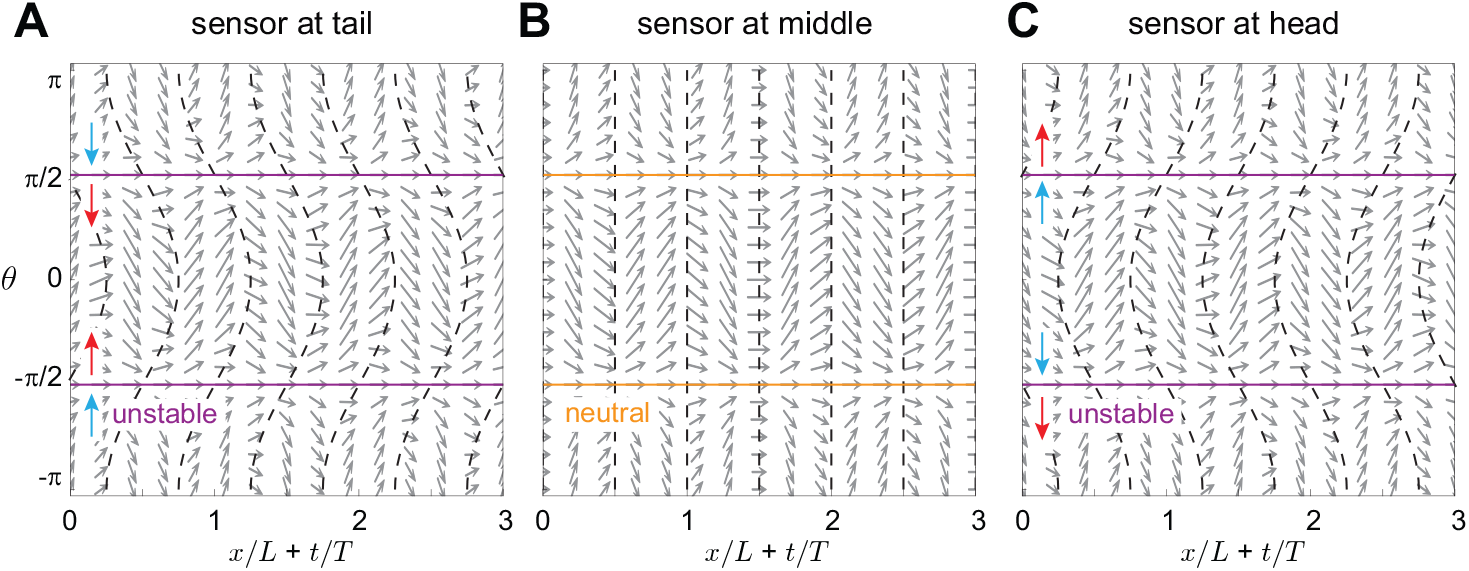
Phase portraits for responding to longitudinal flow gradient. Phase portraits on the phase space (*x* + *λft, θ*) for **A**. sensor at tail (𝓁 = −0.25), **B**. center (𝓁 = 0), and **C**. head (𝓁 = 0.25). Swimming speed *V* = 0.25 and turning rate Ω = 1.

**Figure S5:**
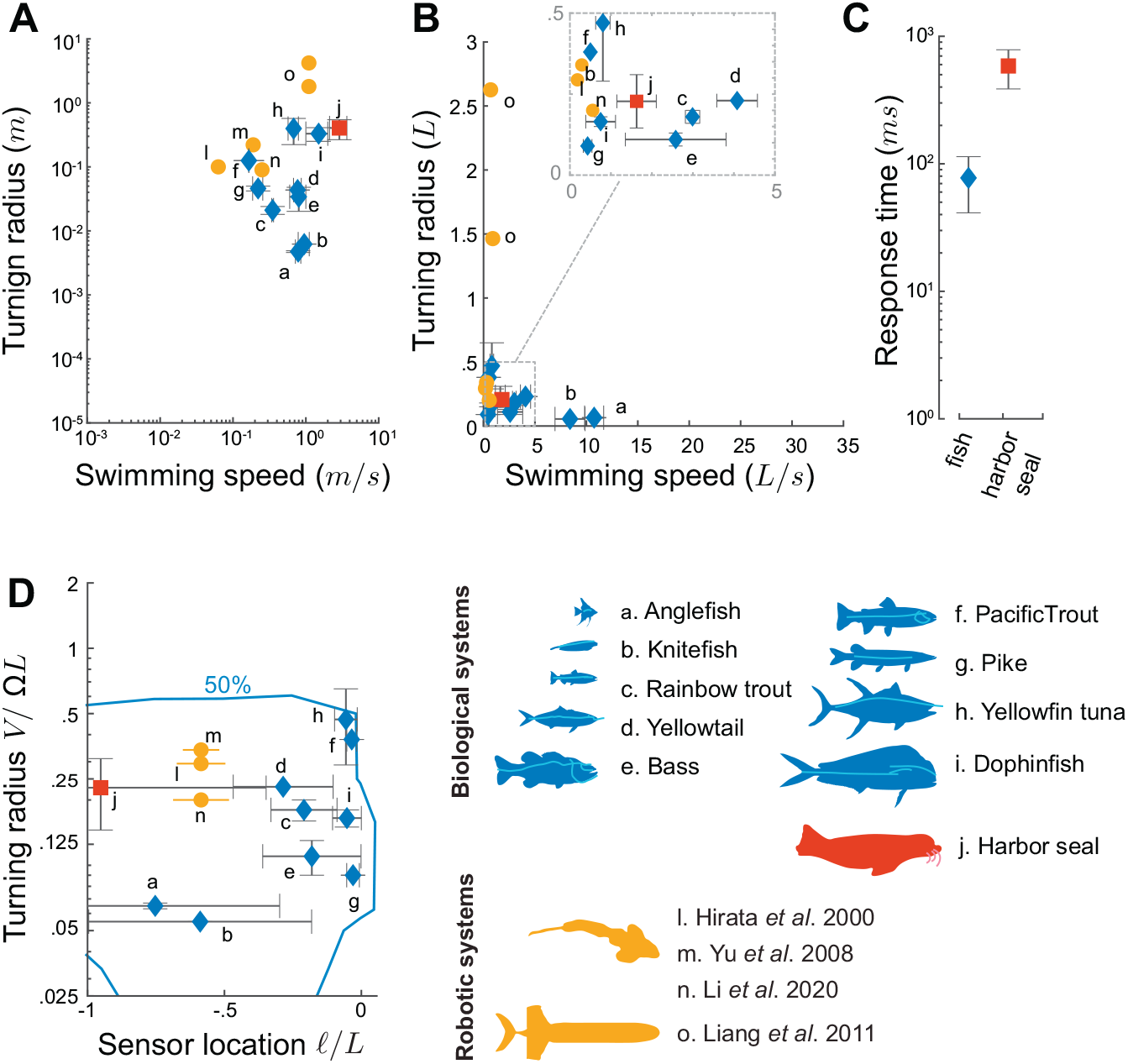
Biological and robotic systems. We gathered biological data from fish, harbor seals and their sea lion relatives, as well as from underwater robotic systems (Table S1). **A**. Turning radius versus swimming speed in log-log scale and dimensional units, and **B**. in units of body length *L*. **C**. Motor response time to mechano-sensory input. Error bars represent the standard deviation of biological data. **D**. Data mapped onto the parameter space (𝓁/*L, V/*Ω*L*) in semi-log scale with sensor location 𝓁 = −*V τ* based on the distance traveled during the neuronal response time *τ*. For robotic fish, we obtained the minimum turning rate from the literature (Table S1) and calculated, based on our model, the range of sensor locations 𝓁/*L* for achieving 95% success rates.

